# A Logical Inquiry of Emotions and Cognition

**DOI:** 10.1101/087874

**Authors:** Arturo Tozzi, Colin James, James F Peters

## Abstract

Experimental procedures in neuroscience rely on the standpoints that the mind is a functional state of the brain and a clear subdivision among different mental faculties does exist in the cortex. According to cognitive neuroscientists, the term "mind" encompasses just the "cognitive" faculties, such as consciousness, perception, thinking, judgement, memory, leaving apart the "emotional" states. Here, taking into account the powerful tools of the first-order predicate logic, we evaluated whether: a) the mind is a function on the physical brain activity; b) different mental faculties can be reduced to a more general one; c) the division of mental faculties in cognition and emotion holds true. We demonstrated that nervous activity is equivalent to mental faculties and that emotions and cognition do not stand for two separated functions of the mind. This means that, counter to our common-sense belief, cognition and emotions are splitted and every faculty of the mind necessarily displays a counterpart in other ones. We point out how it is possible for condensed mind faculties to be unglued in order to become apparently different functions. Therefore, seemingly different mind faculties turn out to be equivalent, because the same logical framework holds for all the types of brain activities, independent of their boundaries and magnitude.

---

The issue of the mind has been tackled by different point of views, from philosophy (Descartes, 1637; Avenarius, 1907; Dennett, 1991) to psychology (Ibáñez et al., 2016) and neuroscience (Başar, 2010). However, there is no universally agreed definition of what its distinguishing properties are. In cognitive neuroscience, the term “mind” refers to a set of “cognitive” faculties (Bosse et al., 2008; Gazzaniga, 2009; Vandekerckhove and Panksepp, 2011). “Cognition” stands here for the mental functions that give rise to information processing: they embrace consciousness, perception, attention, different types of memory, language, learning, thinking, judgement, action, attitudes and interaction in the physical, material, social and cultural world (George, 2003; Almada et al., 2013). In this framework, the emotions (such as joy, fear, love, hate and so on) (Damasio, 2003) are alleged to be primitive and subjective, and therefore are not encompassed in the definition of the mind as such. Other scientists take into account a more general definition of mind, including all mental faculties (Oron Semper et al., 2016). They argue that rational and emotional states cannot be separated, because they are of the same nature and origin, and should therefore be considered all part of it as mind.

There is a general consensus that the mind is correlated with the nervous activity and function of the brain. Therefore, neuroscientists are used to split the brain activity in different subsets of mental faculties, in order that far apart mental domains interact one each other (Touboul, 2012; Gazzaniga, 2013). Indeed, neuroscientific experimental procedures generally aim to assess just specific observational domains of the whole mental faculties (Dricu and Frühholz, 2016). Here we used the powerful tools of the logic, in order to investigate mental functions. Despite scattered skeptical claims (Thagard, 2014), a logical approach has been proven useful in the assessment of mental issues, through modal logic (Hintikka, 1969), partial modal logic (Jaspars and Thijsse, 1996), epistemic logic (Pietarinen, 2003). Further extensions have been used in neuroscience and phenomenology (Pietarinen, 2004). This paper overcomes such uncertainty to clarify the application of mathematical logic to neuroscience, by using the tool of a logic model checker named Meth8. It is based on a variant of Łukasiewicz’ four valued logic system, recently corrected and named VŁ4 (James, 2015; Goodwin, James, 2016; James, 2016). With the help of the first-order predicate logic, our aim was to evaluate whether: a) the mind is a function on the brain; b) mental activities can be reduced to just a general one; c) the above mentioned subdivision of mental faculties in cognition and emotion holds true. Our goal was to explore the possibility that every mental faculty necessarily displays a counterpart in other ones. This would lead to a novel scenario, where different mental faculties were able to scatter, collide and combine, merging together in an assessable way. This logical framework, which holds for all the mental faculties of the brain, independent of their peculiar features, resolution, magnitude and boundaries, aims to assess whether cognitions or emotions are *dual* under logic treatment. The term *dual* refers to a situation where two seemingly different physical systems turn out to be equivalent: if two phenomena are related by a duality, it means that one can be transformed into the other, so that one phenomenon ends up looking just like the other one (Zwiebach and Barton, 2009).

## MATERIALS AND METHODS

### Logic procedure

We mapped categorical subject parts into “literal” variables, through Meth8. We used the 4 propositions < p, q, r, s >. We mapped action relations into respective lists for:

> connectives < And; Not and; Or; Not or; Equivalent; Not equivalent; Imply; Not imply >;
>
> common symbols with ~ for Not as the set {&, ~&, V, ~V, ↔,□, →, ←};
>
> one letter symbols in Meth8 as the set { &, \, +, -, =, @, >, <}.

We also mapped modifiers in modal operators for possibility and necessity, using the following symbols:

> common symbols of the lozenge <> and box [];
>
> one letter characters in Meth8 as % and #.

Predicate logic has two quantifiers, the existential □ as “at least one exists” and the universal ∀ as “for all”. The quantifiers are based on (the misattributed) Aristotle’s Square of Opposition which provably is not bivalent (not exact). However, propositional logic has two modal operators that provably are bivalent (exact) and *generally* interchange with the two quantifiers, <> for □ and [] for ∀.

We also mapped expressions into the format of:

> named types in order of literal, operator, literal such as p & q;
>
> named parts in order of antecedent, connective, consequent.

### Application of logic to mental activity

By using propositional logic, we used the following mapping terminology: p = emotion, q = cognition, r = nervous activity of the brain, and s = mental faculty. Therefore, we can write: <p,q,r,s > = < emotion, cognition, nervous activity, mental faculty >. Following this approach, we tackled two main issues: the mental faculty/nervous activity and the emotion/cognition relationships.

Concerning the first issue, related to mental faculty / nervous activity, we asked whether:

1. Mental faculties are roughly identical with the nervous activity. Format: s=r.
2. Mental faculties are split from nervous activity. Format: s@r.
3. Mental faculties are a functional state of nervous activity. Format: r&s.
4. Mental activities are caused by nervous activity. Format: r>s.
5. Cognition is a functional state of the nervous activity. Format: r&q.
6. Emotion is a functional state of the nervous activity. Format: r&p.
7. Cognition is a mental faculty. Format: q=s.
8. Emotion is a mental faculty. Format: p=s.

Concerning the second issue, related to emotion/cognition relationships, it is possible to determine with logic tools which subdivisions are true, if the two following symbolic assignment approaches can apply:

a. Is it correct to suspect that the most of such subdivisions are not true (not validated), because we assume this using the null hypotheses.
b. We can determine if: emotion and cognition are linked; how much emotion and cognition are linked; emotion and cognition stand for the same activity.

Therefore, we asked whether:

9. Cognition and emotion are exactly the same mental faculty. Format: ((q&s)&(p&s)) = s.
10. Cognition and emotion are two fully split mental faculties. Format: ((q&s)&(p&s)) < s.
11. Cognition and emotion are two linked mental faculties. Format: ((q&s)&(p&s)) > s.

## RESULTS

Concerning the first issue related to mental faculty/nervous activity relationship, we proved that nervous activity is equivalent to mental faculty, as implied by: nervous activity implying emotion but not cognition, and mental faculty implying cognition but not emotion.

Let p = emotion; q = cognition; r = nervous activity; s = mental faculty

If nervous activity implies emotion, and mental faculty implies cognition, and nervous activity does not imply cognition, and mental faculty does not imply emotion, then nervous activity is equivalent to mental faculty.

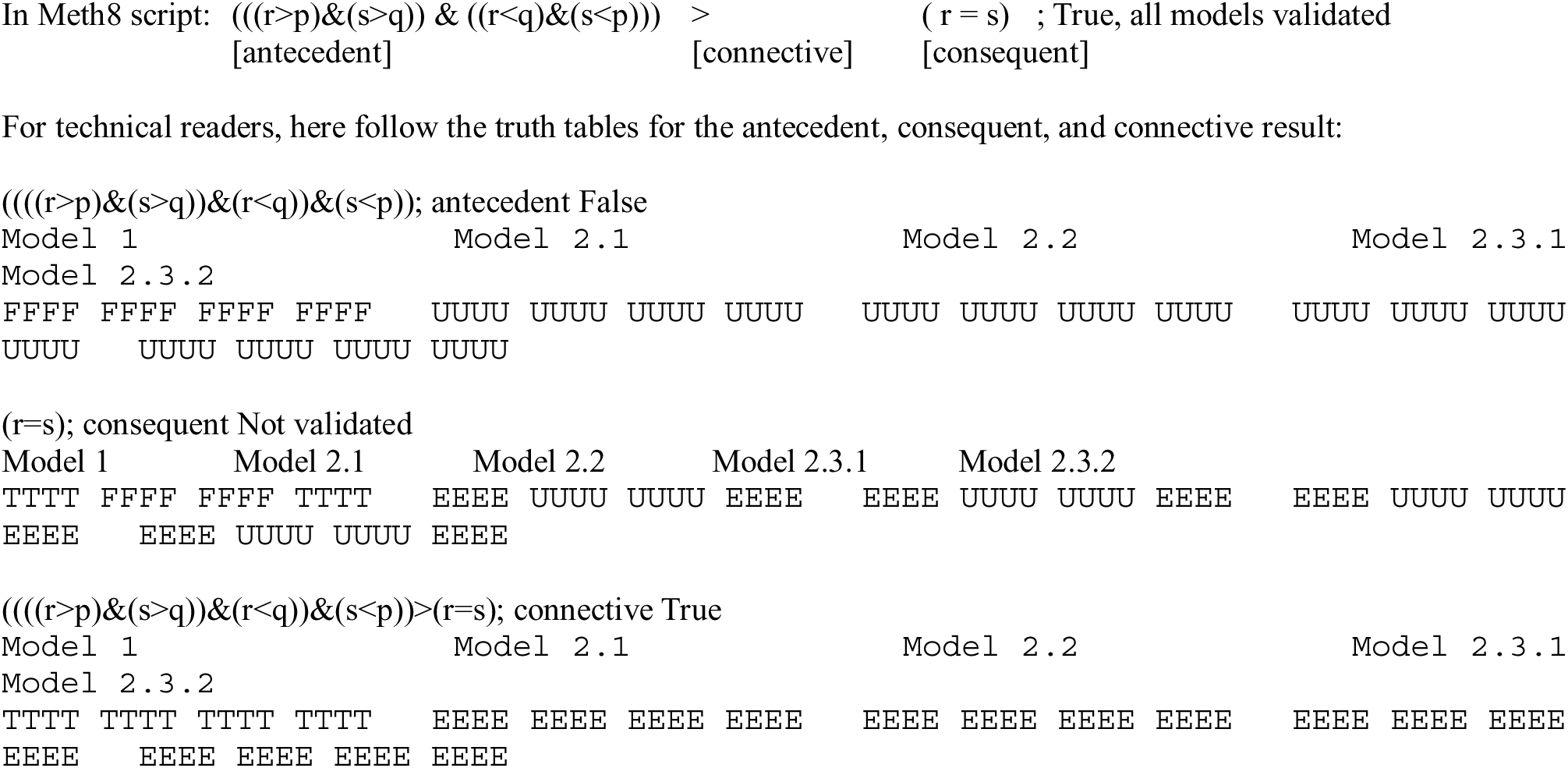

This is an example of a false antecedent implying an invalidated consequent; nevertheless, the expression is a true implication.

Concerning our second issue (emotion/cognition relationships) we achieved the following results:

> Cognition and emotion are exactly the same mental faculty. ((q&s)&(p&s)) = s; Not validated
>
> Cognition and emotion are two fully split mental faculties. ((q&s)&(p&s)) < s; False
>
> Cognition and emotion are two linked mental faculties. ((q&s)&(p&s)) > s; True

The first two expressions, although Not validated, are a good baseline strategy to test the veracity of Meth8, and are included here as a demonstration of that fact. This says that we can produce logical discourse about mental issues. The last, crucial question demonstrates that cognition and emotion are two linked mental faculties.

## DISCUSSION

Here we demonstrate, based on logical tools, that mental faculties are correlated with nervous activity and that cognition and emotion, although believed to be different functions, nevertheless stand for a sole nervous activity. Therefore, our logical approach suggests a topological duality among different mental faculties, because a general nervous activity of the brain might hold for all the types of spatio-temporal nervous functions, independent of their inter- and intra-level relationships, strength, magnitude and boundaries. A question arises: why, by the standpoint of our natural common-sense experience, are we used to split brain activity in different mental faculties? The answer is straightforward, if we take into account topological arguments. If we depict different mental faculties in guise of abstract geometric shapes taking place in the phase space our physical brain, it is possible to demonstrate that they necessarily have at least a feature in common. Various continuous mappings and projections from a cortical zone to the other lead to generalized versions of the Borsuk-Ulam theorem (BUT) (Borsuk 1933;Krantz, 2009; Tozzi and Peters, 2016a), which state that a single shape in a given abstract dimension maps to two identical shapes in one dimension higher (Marsaglia, 1972; Peters, 2016). Brain signals from different mental faculties can be compared, because their two shapes can be assessed at higher-dimensional scales of observation (Tozzi and Peters, 2016b; Peters and Tozzi, 2016a; Weeks, 2002). For technical readers, see also: Matousek (2003).

In the brain, every sub-region encompasses at least one mental faculty, which can be modeled as a shape (Peters and Naimpally, 2012; Di Concilio, 201). Hence, BUT provides a way to evaluate changes of information among different anatomical and functional brain levels in a topological space. Next, consider Brouwer’s fixed point theorem (FPT) (Kellogg et al., 1976). Su (1997) gives a nice illustration of the FPT: no matter how you continuously slosh the coffee around in a coffee cup, some point is always in the same position that it was before the sloshing began. And if you move this point out of its original position, you will eventually we move some other point in the sloshing coffee back into its original position. In BUT terms, this means that not only we can always find a brain region containing an mental faculty, but also that every mental faculty comes together with another one (Peters and Tozzi, 2016b). This means that we can always find a mental faculty, embedded in a cortical area, which is the topological description of another activity, embedded in another area. Therefore, in topological terms, mental faculties/shapes are continually transforming into new equivalent mental faculties/shapes. They might influence each other by scattering, colliding and combining, to create bounded regions in the brain. Eventually mental faculties’ shapes will deform into another, as a result of the collision of a pair of them (Figure). Let a mental faculty be represented in Fig. A. It evolves over time, as it twists and turns through the outer reaches of another mental faculty. An inkling twisting mental faculty appearing in the neighbourhood of the first one is illustrated in Fig. B. The two faculties’ shapes begin interacting, so that the first now has a region of space in common with the second (Fig. C). As a result of the interaction, they become partially stitched together. The partial absorption of one mental faculty into another is shown in Fig. D, where a very large region of the total brain space occupied by the first faculty is absorbed by the second. The two mental faculties become at first concentric in Fig. E., then a complete condensed shape is formed in Fig. F. Therefore, we have the birth of a single condensed mental faculty. This is a further instance of the duality principle in mind theories. That is, one mental faculty is the dual of another, provided the first can be deformed into the second. Hence, it is possible for mental faculties with seemingly varying shapes and sizes to stick together and become a condensed mental faculty. For technical readers, the underlying concepts of homotopy equivalence and Edelsbrunner-Harer nerve are provided in: Peters and Inan (2016) and Peters and Naimpally (2012).

**Figure.**
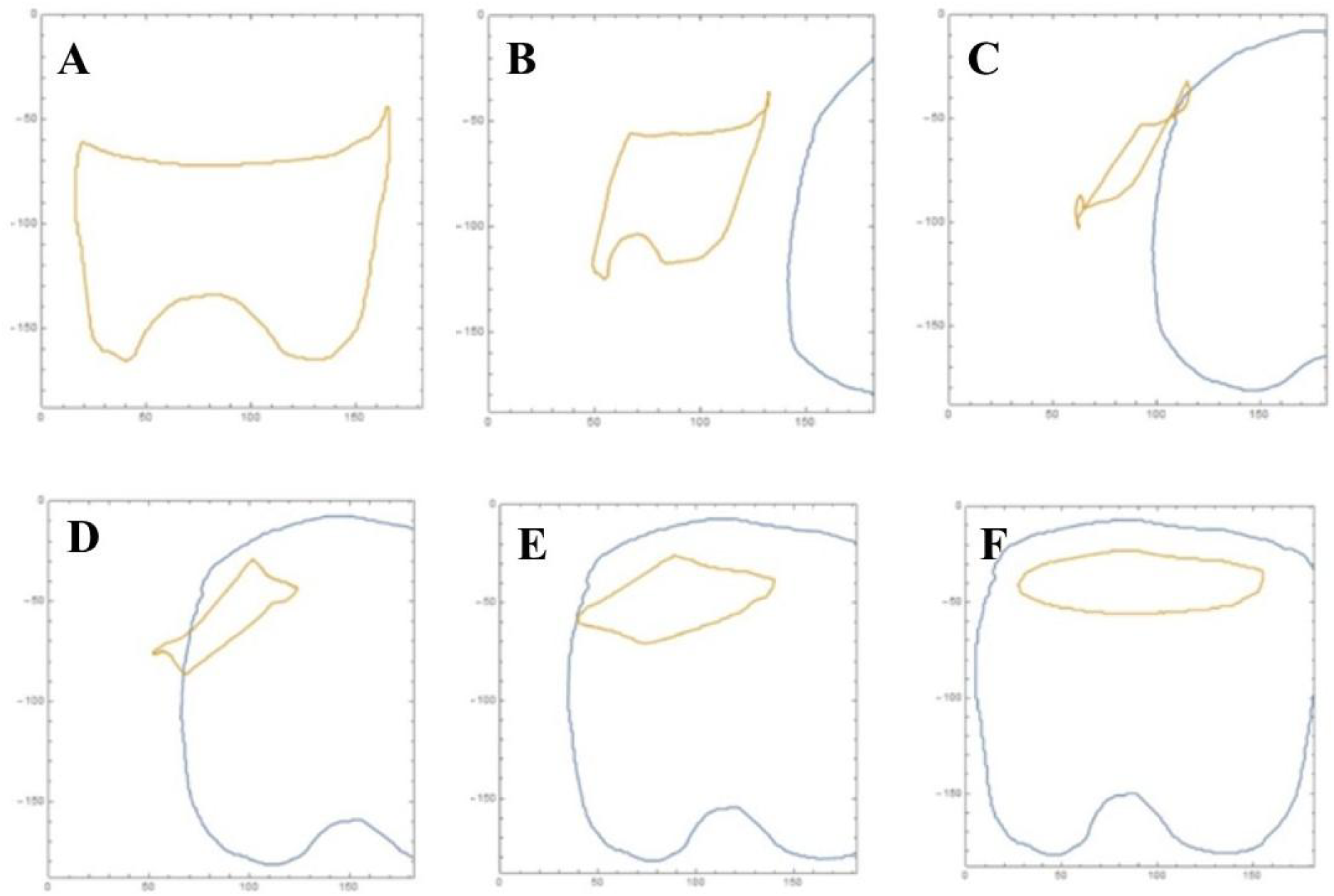
Topological interaction of two shapes, standing for two apparently different mental faculties embedded in the total brain space. **A** depicts a single mental faculty (e.g., emotions). **B**: two mental faculties at two different levels. The second shape stand for cognitive faculties. **C**: interacting mental faculties. **D**: dual mental faculties. **E**: concentric mental faculties. **F**: condensed mental faculty. See text for further details.

Our procedure achieves generalizations that allow the assessment of every possible mental faculty. In other words, there exists an assessable and quantifiable correspondence between the single faculties of the mind. By a common-sense point of view, we are used to conceive mind faculties as too far apart ever to communicate with one another, so that activities bounded on distant brain regions would never have direct contact: for example, two apparently opposite brain activities such as emotions and abstraction have apparently very few in common. However, our topological investigation reveals that this scenario is unfeasible, because there must be at least one element in common also among mental faculties that are apparently very distant one each other. Mental faculties will always have elements in common: every one of them does not exist in isolation, rather they are part of a large interconnected whole. This means that the large repertoire of mind faculties can be described in the same topological fashion. Furthermore, the distinction among different coarse-grained levels of mental faculties does not hold anymore, because different nervous activities observed at small, medium and large neural scales turn out to be topologically equivalent.

## REFERENCES

1) Almada LF, Pereira A Jr, Carrara-Augustenborg C. 2013. What affective neuroscience means for science of consciousness. Mens Sana Monogr. 2013 Jan; 11(1):253–73. doi:10.4103/0973-1229.100409.

2) Avenarius R. 1907. Kritik Der ReinenErfahrung von Richard Avenarius. O.R. Reisland.

3) Başar Erol. 2010. Brain body mind oscillations in scope of uncertainty principle. New York: Springer. p. 5. ISBN 1441961364.

4) Borsuk M. 1933. DreiSatze Uber Die N-Dimensionale EuklidischeSphare. FundamentaMathematicae XX: 177–90.

5) Bosse T, Jonker CM, Treur J. 2008. Formalization of Damasio’s theory of emotion, feeling and core consciousness. Conscious Cogn. 2008 Mar; 17(1):94–113. Epub 2007 Aug 8.

6) Damasio A. 2003. Feelings of emotion and the self. Ann N Y Acad Sci. 2003 Oct; 1001:253–61.

7) Dennett D. 1991. Consciousness Explained. Boston, Massachusetts: Little Brown. ISBN 0-316-18065-3.

8) Descartes R. 1637. Discourse on Method and Meditations on First Philosophy. Hacket Publishing Company, 1998. ISBN 0-87220-421-9.

9) Di Concilio A. 2013. Point-free geometries: Proximities and quasi-metrics, Math. in Comp. Sci. 7, 1, 31–42, MR3043916.

10) Dricu M, Frühholz S. 2016. Perceiving emotional expressions in others: activation likelihood estimation meta-analyses of explicit evaluation, passive perception and incidental perception of emotions. Neurosci Biobehav Rev. 2016 Nov 8. pii: S0149-7634(16)30385-2. doi:10.1016/j.neubiorev.2016.10.020.

11) Gazzaniga MS. 2009. The Cognitive Neurosciences, Fourth Edition. MIT Press. ISBN: 9780262013413

12) Gazzaniga MS. 2013. Shifting gears: seeking new approaches for mind/brain mechanisms. Annu Rev Psychol. 2013; 64:1–20. doi:10.1146/annurev-psych-113011-143817. Epub 2012 Sep 17.

13) George A. 2003. The cognitive revolution: a historical perspective. Trends in Cognitive Sciences 7.

14) Goodwin G, James C. 2016. Meth8 model prover for multivalued logic: Truth as a white light [submitted].

15) Hintikka J. 1969. On the logic of perception. Models for modalities. Dordrecht: Reidel. 151–183.

16) Ibáñez A, García AM, Esteves S, Yoris A, Muñoz E, et al. 2016. Social neuroscience: Undoing the schism between neurology and psychiatry. Soc Neurosci. 2016 Oct 6.

17) James C. 2015. Theorem prover Meth8 applies four valued Boolean logic for modal interpretation. First World Conference: Analogy. Beneméita Universidad Autónoma de Puebla, Mexico, November 4-6, 2015, Handbook, ISBN 978-83-65273-01-1, 50–51.

18) James C. 2016. Language to logic mapper to logic model checker. Verb, clause, and construction Symposium. Universidad de La Rioja.

19) Jaspars J, Thijsse E. 1996. Fundamentals of partial modal logic. Partiality, modality, and non-monotonicity. Stanford: CSLI. 111–141.

20) Kellogg RB, Li TY, Yorke JA. 1976. A constructive proof of the Brouwer fixed point theorem and computational results. SIAM J. Numer. Anal. 13(4): 473–483. doi:10.1137/0713041.

21) Krantz SG. 2009. A Guide to Topology. Edited by The Mathematical Association of America. Washington DC.

22) Marsaglia G. 1972. “Choosing a Point from the Surface of a Sphere.” Annals of Mathematical Statistics 43(2): 645–46. doi:10.1214/aoms/1177692644.

23) Matoušek J. 2003. Using the Borsuk–Ulam Theorem. Lectures on Topological Methods in Combinatorics and Geometry. Berlin Heidelberg: Springer-Verlag.

24) Oron Semper JV, Murillo JI, Bernacer J. 2016. Adolescent Emotional Maturation through Divergent Models of Brain Organization. Front Psychol. 2016 Aug 23; 7:1263. doi:10.3389/fpsyg.2016.01263. eCollection 2016.

25) Peters JF. 2016. Computational Proximity. Excursions in the Topology of Digital Images. Edited by Intelligent Systems Reference Library. Berlin: Springer-Verlag. doi:10.1007/978-3-319-30262-1.

26) Peters JF, Naimpally SA. 2012. Applications of near sets, Notices of the Amer. Math. Soc. 59(4):536–542, DOI:http://dx.doi.org/10.1090/noti817.

27) Peters JF, Inan E. 2016. Strongly proximal Edelsbrunner-Harer nerves. Proc. Jangjeon Math. Soc. 19(3):563–582.

28) Peters JF, Tozzi A. 2016a. Region-Based Borsuk-Ulam Theorem. arXiv.1605.02987

29) Peters JF, Tozzi A. 2016b. String-Based Borsuk-Ulam Theorem. arXiv:1606.04031

30) Pietarinen A-V. 2003. What do epistemic logic and cognitive science have to do with each other? Cognitive Systems Research 4. 169–190.

31) Pietarinen A-V. 2004. Logic, Neuroscience and Phenomenology: In Cahoots? CEUR Workshop Proceedings for WSPI’04.

32) Su FE. 1997. Borsuk-Ulam implies Brouwer: A direct construction. Amer. Math. Monthly 104(9): 855–859, MR1479992.

33) Thagard P. 2014. Cognitive Science. The Stanford Encyclopedia of Philosophy. plato.stanford.edu/entries/cognitive-science/.

34) Tozzi A, Peters JF. 2016a. Towards a Fourth Spatial Dimension of Brain Activity. Cognitive Neurodynamics 10(3): 189–199. doi:10.1007/s11571-016-9379-z.

35) Tozzi A, Peters JF. 2016b. A Topological Approach Unveils System Invariances and Broken Symmetries in the Brain. Journal of Neuroscience Research 94(5): 351–65. doi:10.1002/jnr.23720.

36) Vandekerckhove M, Panksepp J. 2011. A neurocognitive theory of higher mental emergence: from anoetic affective experiences to noetic knowledge and autonoetic awareness. Neurosci Biobehav Rev. 2011 Oct; 35(9):2017–25. doi:10.1016/j.neubiorev.2011.04.001. Epub 2011 Apr 19.

37) Weeks JR. 2002. The shape of space, IInd edition. Marcel Dekker, inc. New York-Basel.

38) Zwiebach, Barton. 2009. A First Course in String Theory. Cambridge University Press. ISBN 978-0-521-88032-9.

